## Supplementary Figures for "Evaluating MINFLUX experimental performance *in silico*"

### Supplementary Information

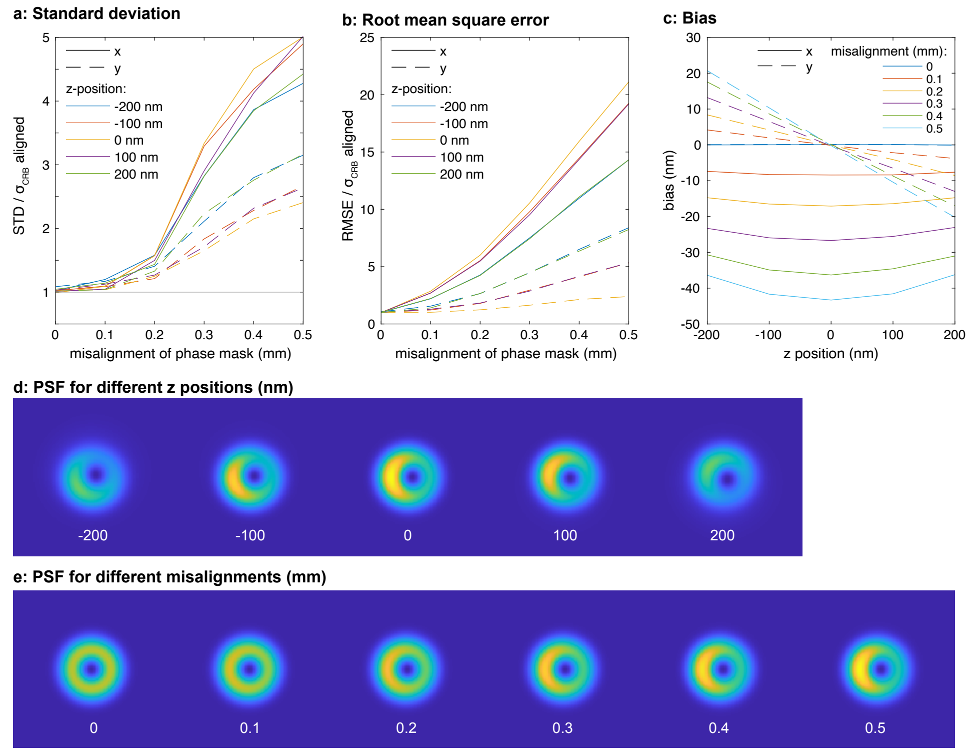


**Supplementary Information Fig. 1. Misalignment of the vortex phase pattern**. Simulation of how a misalignment of a vortex phase pattern along the x-direction affects fluorophore localization. **a,** The ratio of the standard deviation (STD, see Data Analysis section of Methods) of the fluorophore's position estimate to the Cramér-Rao Bound (CRB) for a perfectly aligned phase pattern as a function of misalignment, investigated for fluorophores at varying distances from the focal plane. Misalignment has the greatest effect on STD in the direction of misalignment. **b,** The normalized root mean square error (RMSE, see Methods) of the true and estimated fluorophore position as a function of misalignment. **c,** Bias (see Methods) of the estimated fluorophore position as a function of the fluorophore's distance from the focal plane. **d,** The point spread function (PSF) at different z positions. **e,** The PSF at the plane of focus (z=0) for misalignments in steps of 0.1 mm. For all plots, L=75 nm, vortex phase mask, a scan pattern consisting of 6 orbit plots plus their centroid, point dwell time of 0.1 ms, average photons (aligned case) of 45 per localization, iterative position estimator. See example 2c (example2c_phaseplate_misalignment.m or .ipynb).

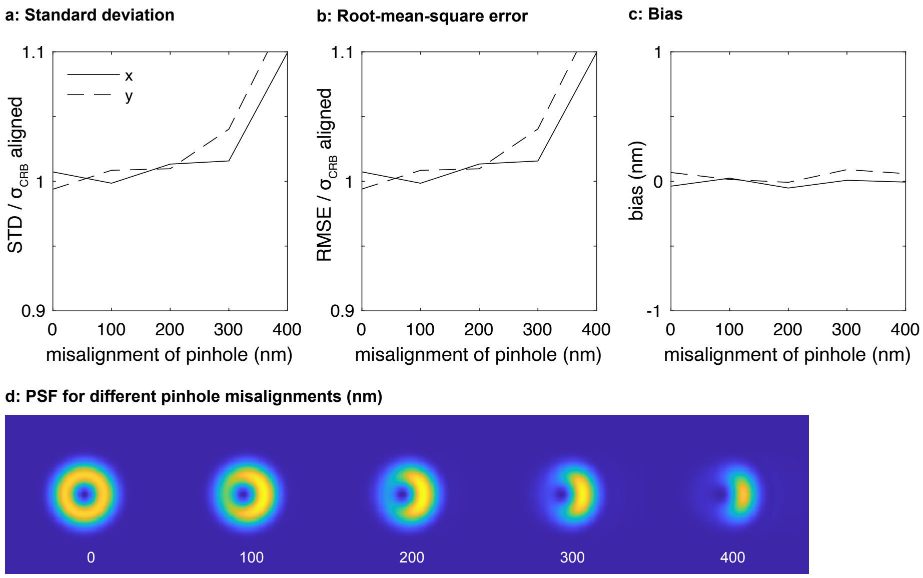


**Supplementary Information Fig. 2. Misalignment of the pinhole.** Simulation of how a misalignment of the pinhole along the x-direction affects fluorophore localization. **a,** The ratio of the standard deviation (STD, see Data Analysis section of Methods) of the fluorophore's position estimate to the Cramér-Rao Bound (CRB) for a perfectly aligned pinhole as a function of misalignment. Misalignment has limited effect on STD. **b,** The normalized root mean square error (RMSE, see Methods) of the true and estimated fluorophore position as a function of pinhole misalignment. Misalignment has limited effect on RMSE. **c,** Bias (see Methods) of the estimated fluorophore position as a function of pinhole misalignment. There is no effect. **d,** The PSF at the plane of focus (z=0) for pinhole misalignments in steps of 100 nm. For all plots, L=75 nm, vortex phase mask, a scan pattern consisting of 6 orbit plots plus their centroid, point dwell time of 0.1 ms, average photons (aligned case) of 72, iterative position estimator. See example 2d (example2d_pinhole_misalignment.m or .ipynb).

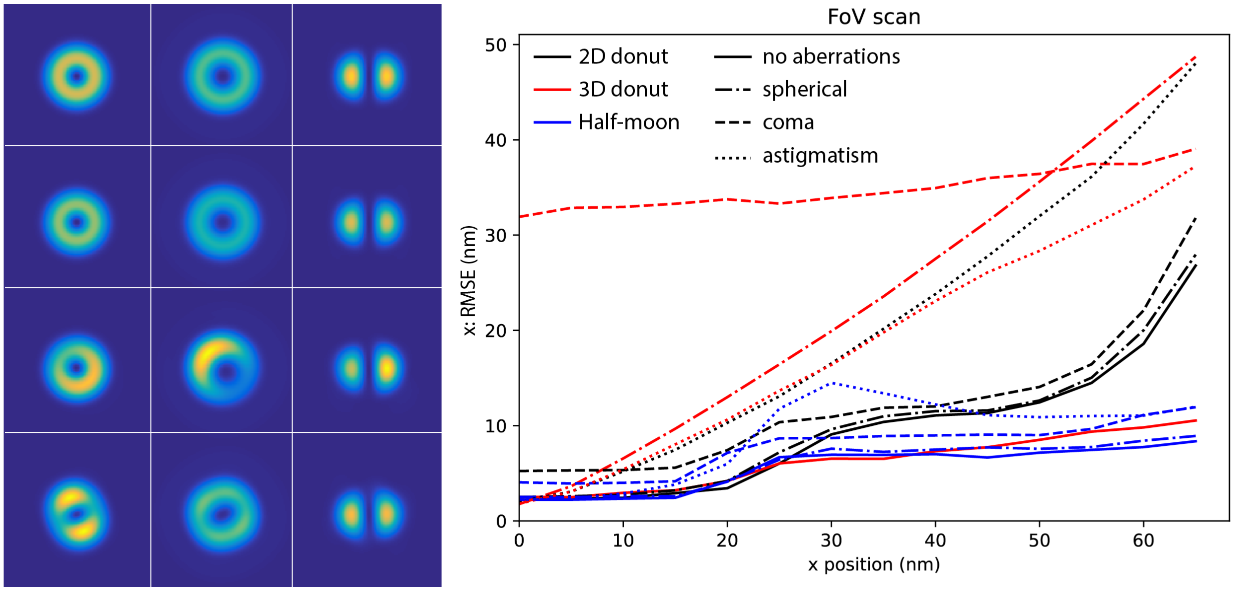


**Supplementary Information Fig. 3. Aberrations.** Effects of spherical, coma, and astigmatism aberrations on 2D and 3D (top hat phase mask) donut (vortex phase mask) and phase flux (halfmoon phase mask) point spread functions. **Left,** shown in the plane of focus. **Right**, RMSE (see Methods) of position estimate and true position after scanning each PSF along a single dimension at -L/2, 0, L/2 with L=75 nm, fluorophore brightness = 1000, point dwell time = 0.1 ms, 5000 localizations. All aberrations applied are 0.15 rad. In the case of coma and astigmatism, this is split as a sum of 0.075 rad each along the vertical and horizontal directions in the pupil plane. See example 2b (example2b_compare_PSFs.m or .ipynb).

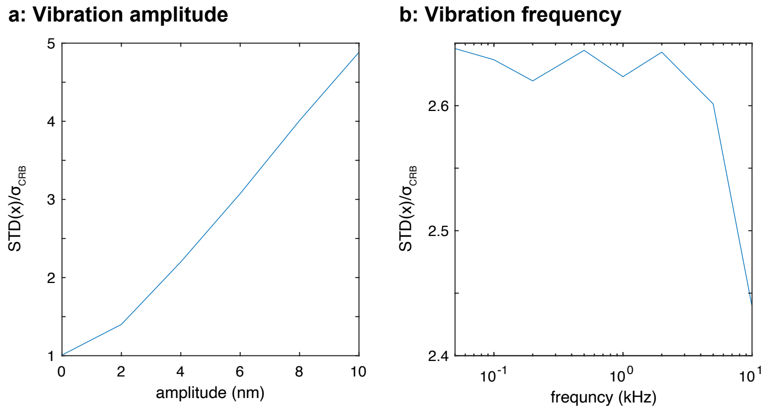


**Supplementary Information Fig. 4. System Vibrations.** The effect of system vibrations on standard deviation of fluorophore position estimate (STD, see methods). Vibrations were simulated using moving fluorophores (“FlMoving” class, see Methods). **a,** Fluorophores oscillating at a frequency of 100 Hz with varying amplitudes. Increased amplitude leads to decreased precision in fluorophore position. **b,** Fluorophores moving at varying frequences with an amplitude of 5 nm. STD has low dependence on frequency. In all experiments, fluorophores were localized with the 2D tracking sequence, Tracking_2D.json. See example 5 (example5_moving_fluorophore.m or .ipynb).

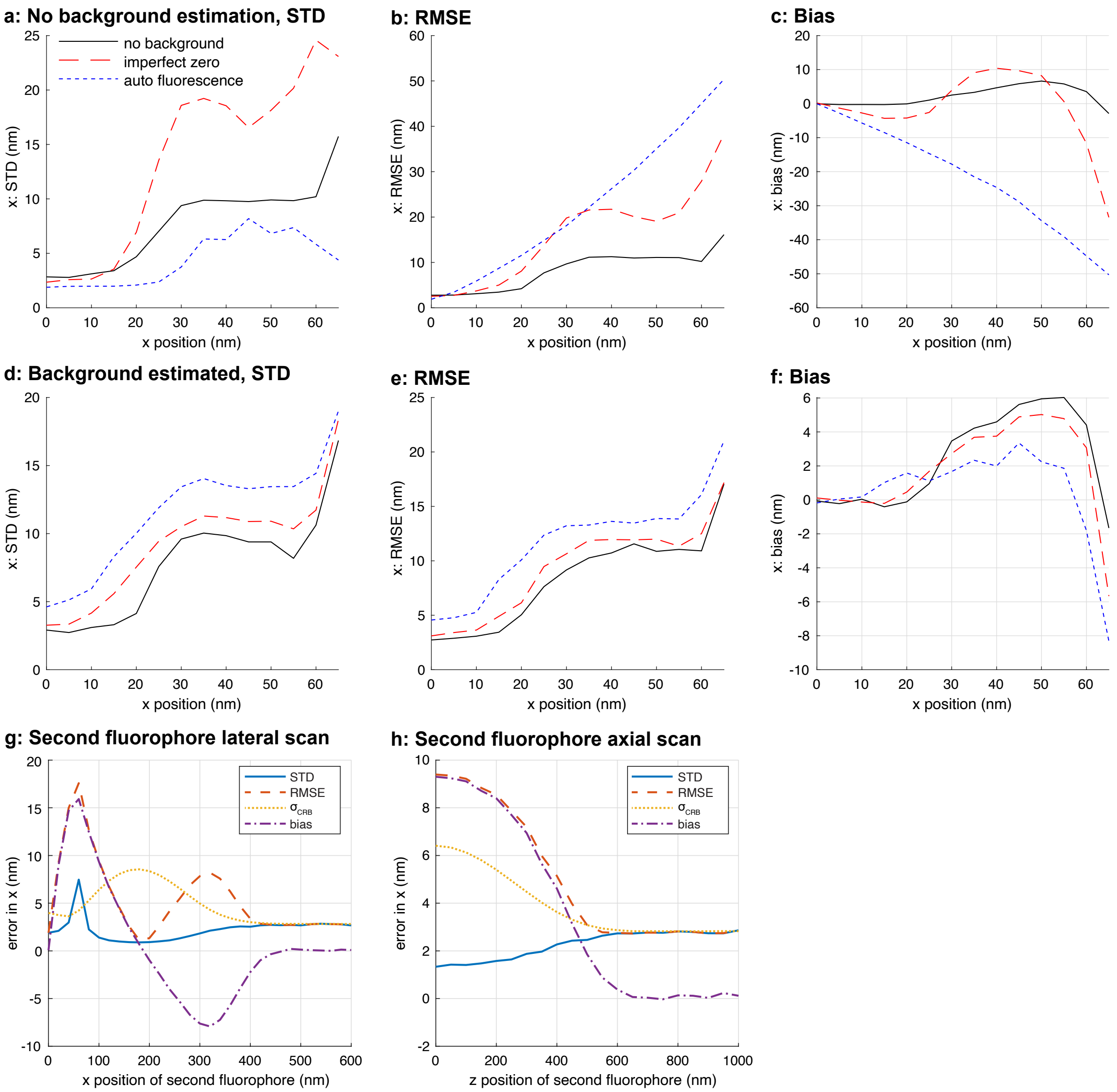


**Supplementary Information Fig. 5. Background.** Effects of background from imperfect zero, from autofluorescence, and from nearby fluorophores on emitter localization. **a-c,** The background is not considered, which leads to a bias in the iterative quadratic LSQ estimator. **d-f,** The background is estimated and subtracted from the photon counts before estimation, which removes the bias. The background still causes an increased localization error (STD). **g-h,** Effect of a nearby fluorophore of equal brightness.

Background values: Zero-offset (psf.zerooffset): 0.01; auto fluorescence background (sim.background): 30 kHz. L=75 nm. Position of the target fluorophore: [0,0,0]. Position of nearby fluorophore at [50 50 400] nm unless otherwise indicated. 100 photons/localization (no background case). See example 8 (example8_background.m or .ipynb).

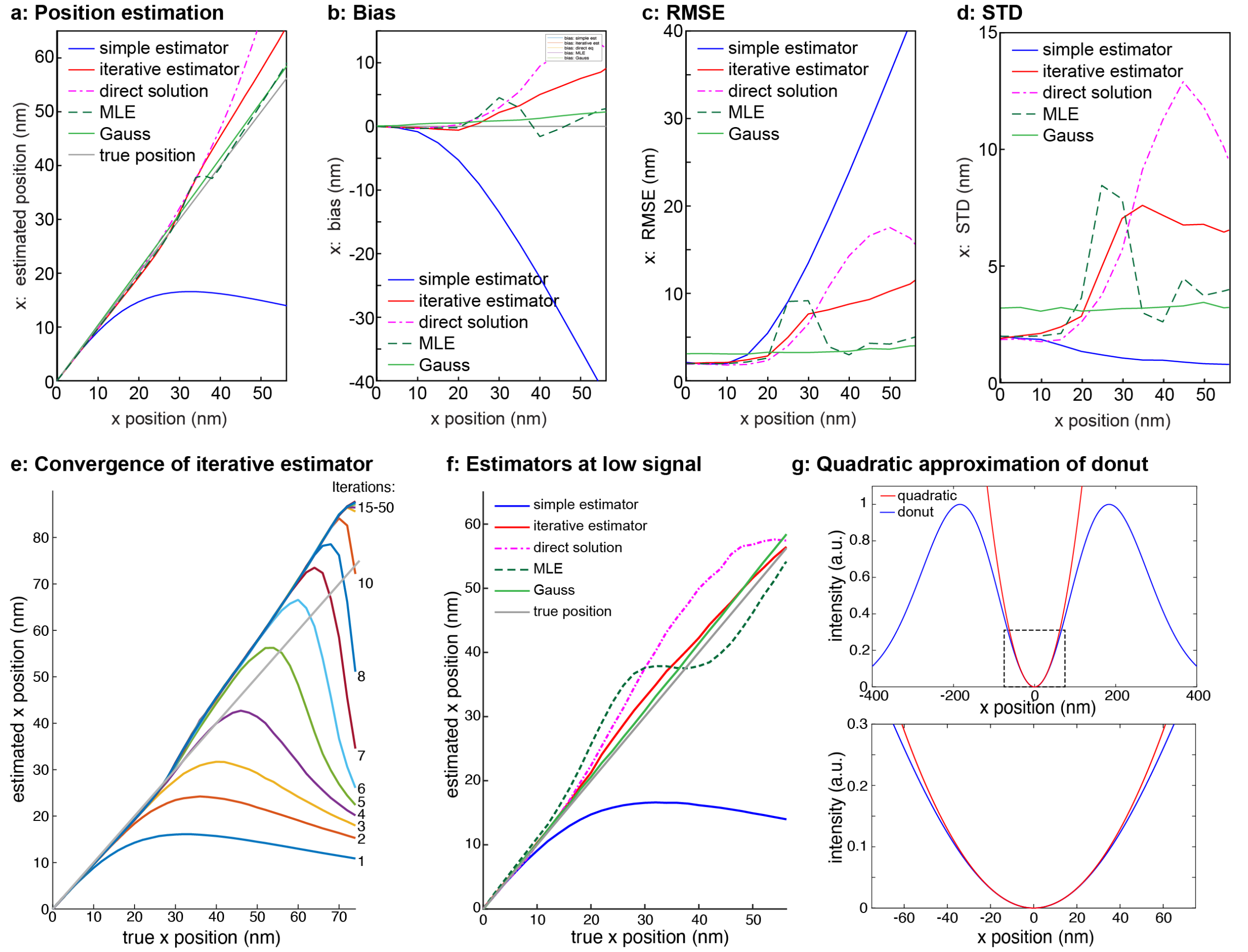


**Supplementary Information Fig. 6. Estimators. a, Position, b, Bias, c, RMSE and d, STD** compared for the estimators used in this manuscript. Note that a strong bias, as for the simple estimator, can lead to low, but meaningless STD. **e,** Convergence of the iterative estimator with a quadratic approximation (Methods, Eq. 16) over repeated scans of a fluorophore with a donut-shaped point spread function. As the number of iterations increases, the estimated x position converges to the true estimated position over a larger field of view, with some bias after 30 nm due to mismatches between the quadratic approximation of the excitation PSF and the donut excitation, shown in **g**. **f,** The performance of various estimators, described in Methods, with only 15 signal photons collected. The iterative estimator and the Gaussian approximation via least squares perform best in these low light level conditions. Notably, the maximum likelihood estimate (MLE) has the strongest bias for low photon counts. **g,** A plot of a donut shape and its quadratic approximation. Note that the quadratic approximation only matches the donut near x=0 nm. Bottom panel shows the black dashed box zoomed in for detailed comparison. Estimators using a quadratic approximation of a donut fail when the fluorophore is not near the donut center.

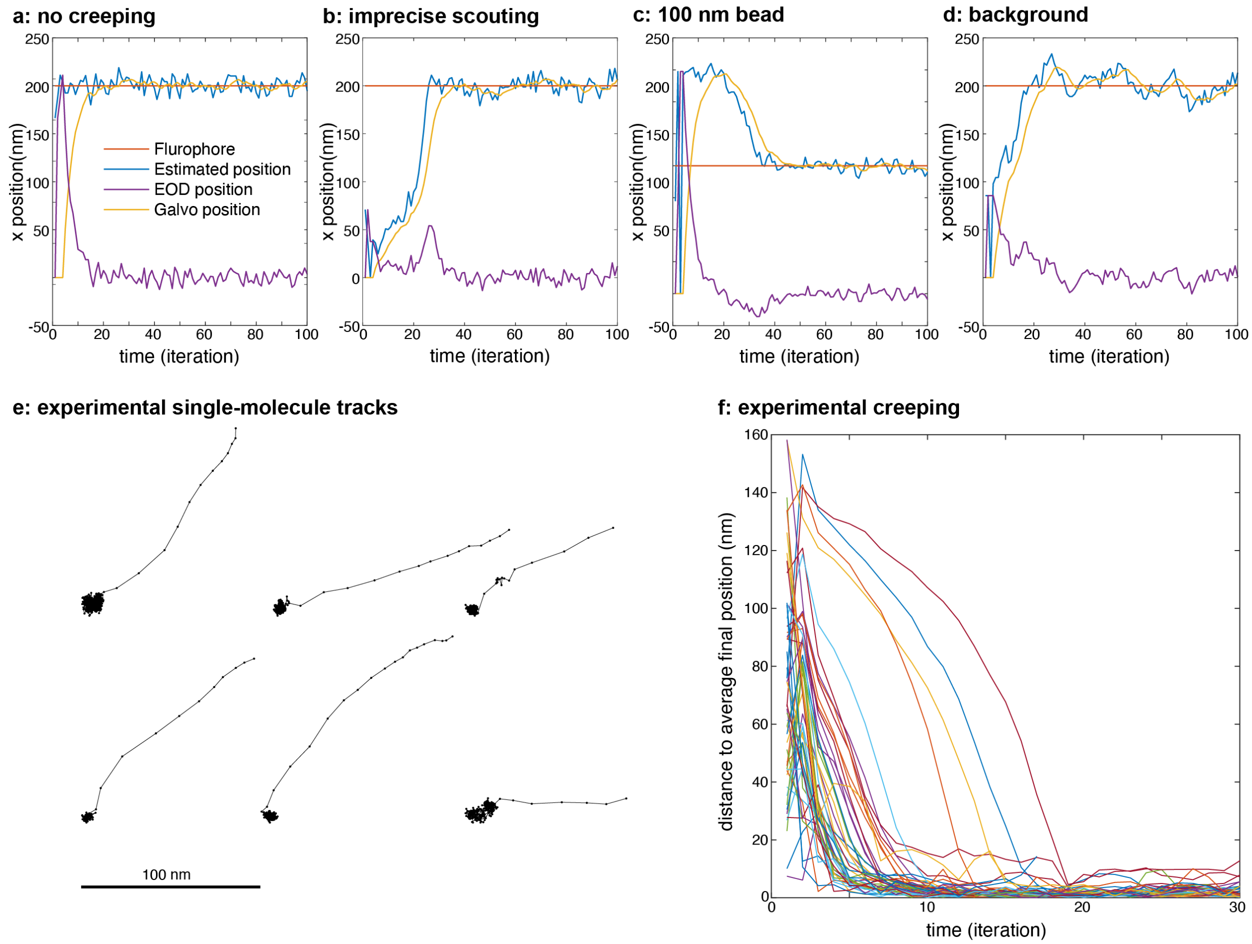


**Supplementary Information Fig. 7. Creeping of estimated fluorophore position. a,** Traces of estimated position, true fluorophore position, galvo (or piezo, slow scanner) position, and electro-optical deflector (EOD) position in an ideal case. The estimated position quickly converges to the true fluorophore position. **b,** In the case of imprecise scouting (a Gaussian estimator with $\sigma = 80$ nm), our initial guess is far from the true fluorophore position, and often it takes many iterations to converge. **c,** In the case of localizing a 100 nm bead, the size of the bead causes a blurring of the PSF and increases the intensity at the center, which leads to a bias in the estimator and slow convergence of the estimator to the centroid position of the bead. **d,** In the case of high background (30 kHz), the resulting bias in the estimator will lead to a similar slow convergence. Shown are instructive example tracks. Creeping is stochastic with a high variability, thus even in case of imperfections, a fraction of tracks converged quickly.

In all simulations, a fluorophore was placed at (200, 50, 0) nm and was localized with the 2D tracking sequence, Tracking_2D.json. Simple 2D Gaussian (see Methods Eq. 13 Linearized Least Squares Estimator) and donut (see Methods Eq. 8 Balzarotti et al.[1], Eq. S50) estimators were used. See example 3 (example3_Abberior_sequence.m or .ipynb).

**e,** Experimental example tracks of single fluorophores (Atto647N) immobilized on a coverslip show creeping at the beginning of the tracks. **f,** Magnitude of creeping vs. iteration for 50 single molecules. Only valid final iterations are plotted.

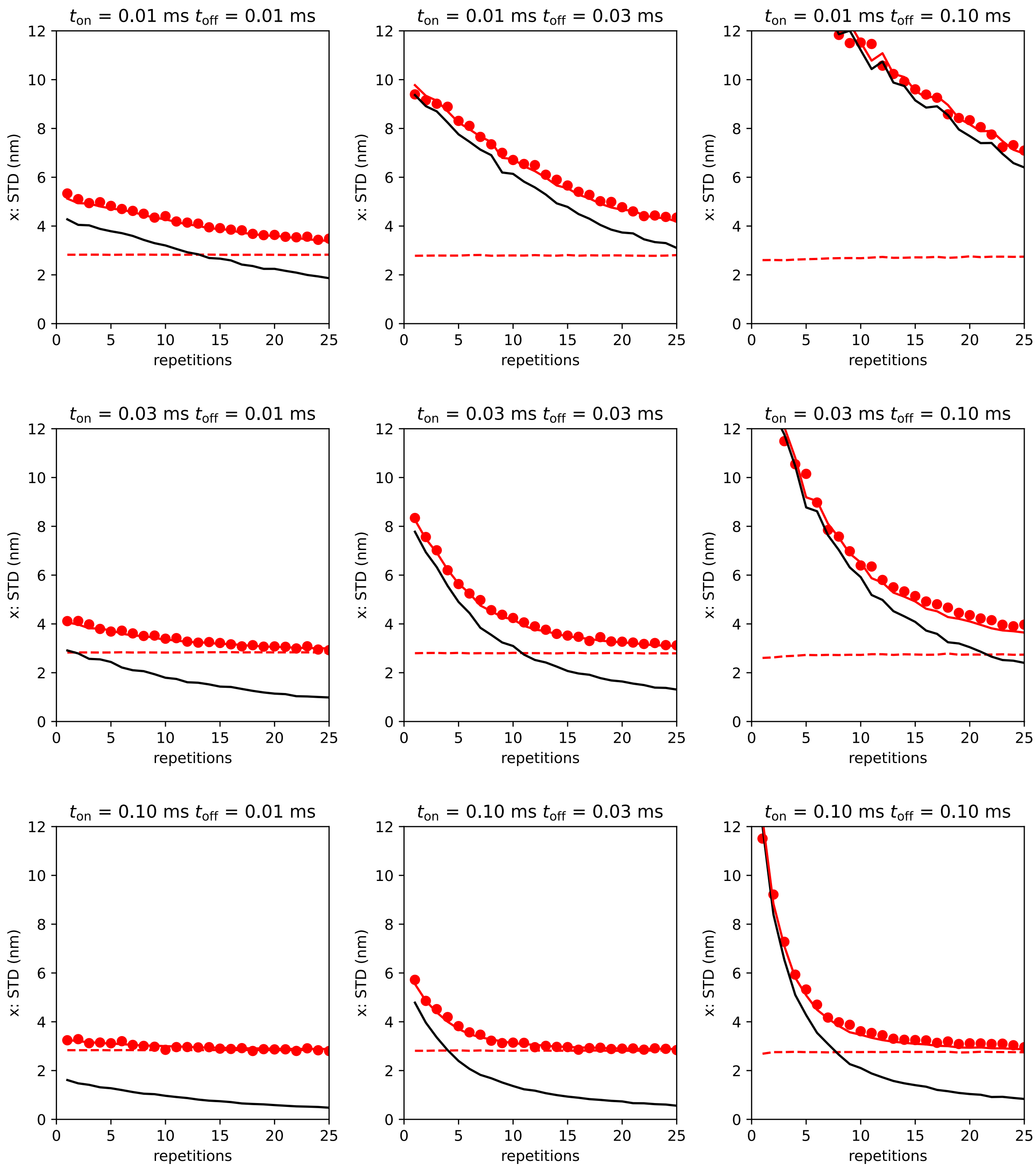


**Supplementary Information Fig. 8. Loss in precision caused by fluorophore flickering, effect of fluorophore on and off time and microscope dwell time.** Here we plot the standard deviation of a localization estimate vs the number of repetitions of a single flickering fluorophore with varying on and off times. The pattern dwell time is fixed to 400 μs and the total number of photons is set at 100. Red dots represent the measured standard deviation of a fluorophore position. The red dashed line is the CRB for these measurements. The black line is the extra error for very high photon counts. The red line $\mathrm{STD}(r, x)$ is calculated from the CRB and the extra error as described in the main text, for 100 photons.
See example 4 (example4_blinking_fluorophore.m or .ipynb).

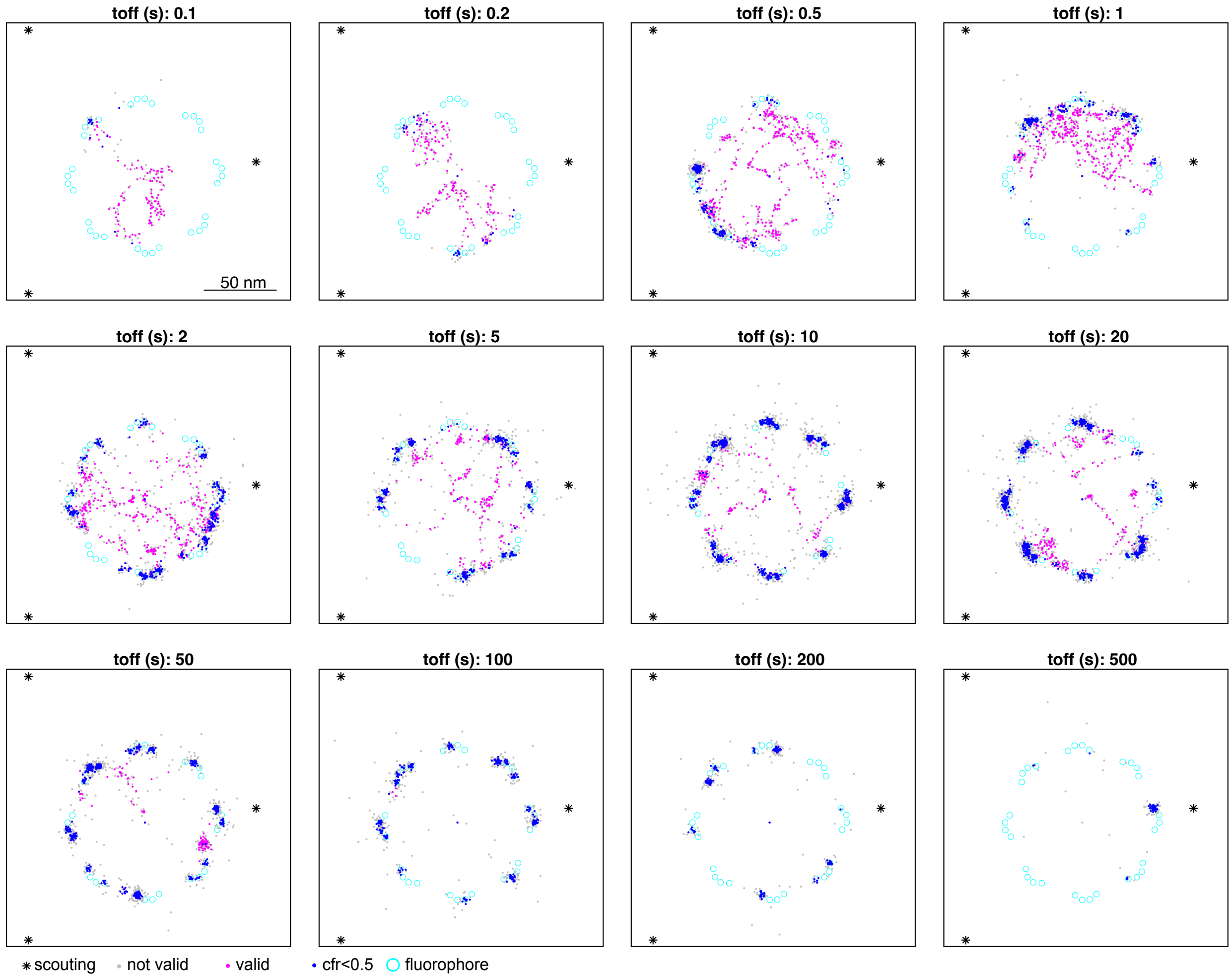


**Supplementary Information Fig. 9. Optimizing fluorophore density.** Effects of the off time of the dSTORM probe on localization density. Short off times result in many fluorophores being on at once, which means the sparsity requirement is not met. This leads to wrong localizations and prevents the CFR check (see Methods, Abberior MINFLUX model) from passing. Long off times results in most fluorophores being off when scouting or localization occurs and so they are never seen in the limited measurement time. For all experiments, we used a photon budget 5000, 2 reactivations, an Abberior 2D imaging sequence, Imaging_2D.json, and a measurement time of 60 s. See example 10 (example10_imaging.m or .ipynb).

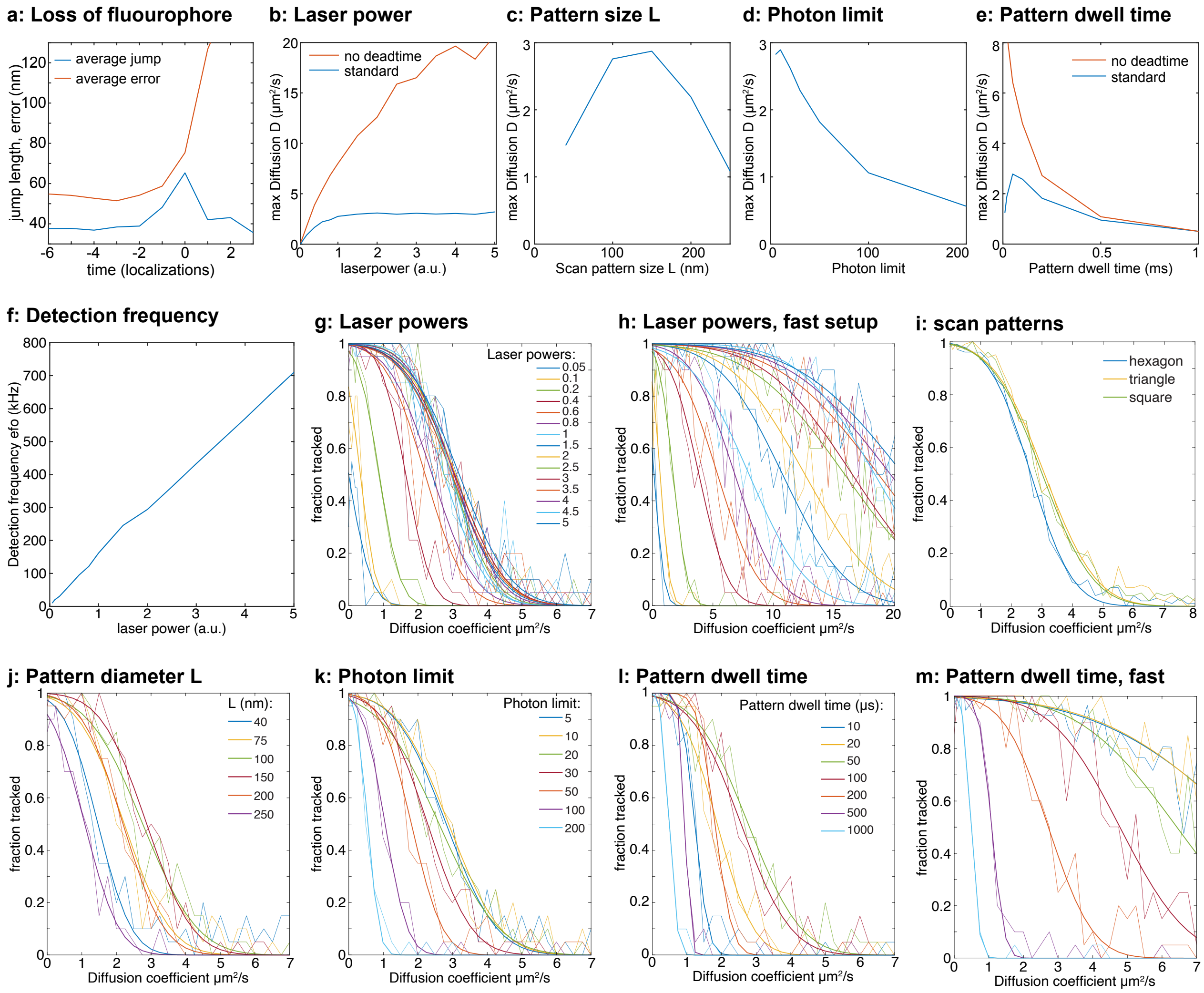


**Supplementary Information Fig. 10. Optimizing a MINFLUX tracking experiment in silico. a,** Jump length and localization error aligned by the last tracked localization and averaged over 300 tracks, shows that tracking fails after large random displacements. **b-e**, Systematic analysis of the maximum diffusion coefficient $D_{\max}$ (µm/s$^2$) that can be measured for different laser powers, pattern sizes, photon limits and pattern dwell times for the standard Abberior MINFLUX and for an equivalent microscope with no dead times. $D_{\max}$ increases with the laser power, but levels off quickly because of the limited time resolution of standard instruments. Note that high laser powers lead to high count rates of the fluorophores (**f**), which might become unrealistic. **g-m**, Fraction of successfully tracked molecules (defined as >90 consecutive localizations with an error below 100 nm) vs. the diffusion coefficient and sigmoidal fit. For each diffusion coefficient 20 tracks were simulated.

After choosing optimal values based on these parameter scans and optimal scan patterns, while keeping the same fluorophore emission rates, we could improve $D_{\max}$ from 2.5 µm$^2$/s to 4.2 µm$^2$/s (see Figure 2l, example 11, example11_max_diffusion.m or .ipynb).

| Reference | Masullo [18] | Masullo [13] | He [16] | Slenders [17] | Srambickal [19] | Rosati [20] | Liu [14] | This work |
|---|---|---|---|---|---|---|---|---|
| Code available | ✓ | ✓ | ✗ | ✓ | ✗ | ✗ | ✓ | ✓ |
| Vectorial PSF model | ✗ | ✗ | ✓ | ✓ | ! | ✗ | ✓ | ✓ |
| 3D MINFLUX | ✗ | ✗ | ✗ | ✗ | ✗ | ✗ | ✗ | ✓ |
| Pinhole detection | ✗ | ✗ | ✗ | ✓ | ✗ | ✗ | ✗ | ✓ |
| Scouting, imaging of fluoropore patterns | ✗ | ✗ | ✗ | ✗ | ✗ | ✗ | ✗ | ✓ |
| Pattern scan with different PSFs | ✗ | ✗ | ✗ | ✗ | ✗ | ✗ | ✓ | ✓ |
| Estimators | ✓ | ✗ | ✓ | ✗ | ✓ | ✓ | ✗ | ✓ |
| Iterative MINFLUX with pattern recentering | ✗ | ✗ | ✗ | ✗ | ✓ | ✗ | ✗ | ✓ |
| Realistic hardware timing | ✓ | ✗ | ✗ | ✗ | ✓ | ✗ | ✗ | ✓ |
| Fluorophore Blinking | ✓ | ✗ | ✗ | ✗ | ✓ | ✗ | ✗ | ✓ |
| Fluorophore Bleaching | ✗ | ✗ | ✗ | ✗ | ✓ | ✓ | ✗ | ✓ |
| Fluorophore Movement | ✗ | ✗ | ✗ | ✗ | ✗ | ✗ | ✗ | ✓ |
| Multiple Fluorophores | ✗ | ✗ | ✗ | ✗ | ✗ | ✗ | ✗ | ✓ |
| CRB calculations | ✓ | ✓ | ✓ | ✓ | ✗ | ✓ | ✓ | ✓ |
| RMSE/Bias | ✓ | ✗ | ✓ | ✗ | ✗ | ✓ | ✗ | ✓ |
| Abberior sequence files | ✗ | ✗ | ✗ | ✗ | ✗ | ✗ | ✗ | ✓ |

| | |
|---|---|
| Yes | ✓ |
| Unclear | ! |
| No | ✗ |

**Supplementary Information Table 1. Comparison of available MINFLUX simulations.**


18. Masullo, L. A. *et al.* Pulsed Interleaved MINFLUX. *Nano Lett.* **21**, 840–846 (2021).
13. Masullo, L. A., Lopez, L. F. & Stefani, F. D. A common framework for single-molecule localization using sequential structured illumination. *Biophys. Rep.* **2**, 100036 (2022).
16. He, C. *et al.* Effects of optical aberrations on localization of MINFLUX super-resolution microscopy. *Opt. Express* **30**, 46849–46860 (2022).
17. Slenders, E. & Vicidomini, G. ISM-FLUX: MINFLUX with an array detector. *Phys Rev Res* **5**, 023033 (2023).
19. Srambickal, C. V. *et al.* Near-infrared MINFLUX imaging enabled by suppression of fluorophore blinking. *bioRxiv* 2024.08.27.609859 (2024) doi:10.1101/2024.08.27.609859.
20. Rosati, M., Parisi, M., Gianani, I., Barbieri, M. & Cincotti, G. Fundamental precision limits of fluorescence microscopy: a perspective on MINFLUX. *Opt. Lett.* **49**, 4938–4941 (2024).
14. Liu, Y., Dong, J., Augusto Maya, J., Balzarotti, F. & Unser, M. Point-spread-function engineering in MINFLUX: optimality of donut and half-moon excitation patterns. *Opt. Lett.* **50**, 37–40 (2025).


| Parameter | Default Value | Optimized Value |
|---|---|---|
| **patGeoFactor** | 0.28 | 0.4 |
| **phtLimit (count)** | 20 | 10 |
| **patDwellTime (μs)** | 100 | 50 |
| **pattern** | hexagon | square |

**Supplementary Information Table 2. Optimized values for last MINFLUX iteration in diffusion experiment.** See example 11, example11_max_diffusion.m or .ipynb.
