## Supplementary Mathematica notebook (.nb) for "Evaluating MINFLUX experimental performance *in silico*": SimuFLUX_estimators.pdf

### Estimators used in SimulFLUX

#### Equations

Calculate normalized probabilities from PSF:

In[ ]:=
```
pnorm[fx_, coordsx_] :=
 Assuming[{L > 0, σq > 0}, With[{px = Map[Simplify[Apply[fx]], coordsx]},
   FullSimplify[px / FullSimplify[Total[px]]]]]
```

Solution to the leased squares problem from Jacobian:

In[ ]:=
```
lsq[J_, Y_] := Assuming[{L > 0, σq > 0, x0 ∈ Reals, y0 ∈ Reals},
  With[{JT = Transpose[J]}, Simplify[Dot[Inverse[Dot[JT, J]], Dot[JT, Y]]]]]
```

Quadratic approximation of donut:

In[ ]:=

$$f_q[x\_] := A \frac{(x - x0)^2}{2 * \sigma^2} + A * b$$

In[ ]:=
```
ang2x[tx_, R_] := R * Cos[tx]
ang2y[ty_, R_] := R * Sin[ty]
```

#### LSQ for 1D quadratic approximation

Experiment : measure ni. Experimental probabilities pi=ni/sum(ni)
Define L2=L/2;

##### At x0=0, 2 measurements

Linearization of p_i. From Balzarotti 2017, eq. xxx

In[ ]:=
```
pn = pnorm[fq, {{-L2}, {L2}}];
J = FullSimplify[D[pn, {{x0}}] /. x0 → 0];
pv = {p1, p2};
Y = pv - pn /. x0 → 0 // Simplify;
r = lsq[J, Y]
```

Out[ ]=

$$\left\{\frac{(p1 - p2)\left(L2^2 + 2\, b\, \sigma^2\right)}{2\, L2}\right\}$$

#### At arbitrary x0, for iterative LSQ

In[ ]:=
```
pn = pnorm[f_q, {{-L2}, {L2}}];
J = FullSimplify[D[pn, {{x0}}]];
Y = pv - pn // Simplify;
r2 = lsq[J, Y]
```

Out[ ]=

$$\left\{\frac{\left(L2^2 + x0^2 + 2\,b\,\sigma^2\right)\left(L2^2\,(p1 - p2) - 2\,L2\,x0 + (p1 - p2)\left(x0^2 + 2\,b\,\sigma^2\right)\right)}{2\,L2\left(L2^2 - x0^2 + 2\,b\,\sigma^2\right)}\right\}$$

replace probabilities by difference: d12=p1-p2

In[ ]:=
```
r2r = r2 /. p1 → d12 + p2 // FullSimplify
```

Out[ ]=

$$\left\{\frac{\left(L2^2 + x0^2 + 2\,b\,\sigma^2\right)\left(-2\,L2\,x0 + d12\left(L2^2 + x0^2 + 2\,b\,\sigma^2\right)\right)}{2\,L2\left(L2^2 - x0^2 + 2\,b\,\sigma^2\right)}\right\}$$

no background:

In[ ]:=
```
r2r /. b → 0
```

Out[ ]=

$$\left\{\frac{\left(L2^2 + x0^2\right)\left(-2\,L2\,x0 + d12\left(L2^2 + x0^2\right)\right)}{2\,L2\left(L2^2 - x0^2\right)}\right\}$$

#### At arbitrary x0 with center measurement

In[ ]:=
```
pn = pnorm[f_q, {{-L2}, {L2}, {0}}];
J = FullSimplify[D[pn, {{x0}}]];
pv = {p1, p2, p0};
Y = pv - pn // Simplify;
rc = lsq[J, Y] // FullSimplify
```

Out[ ]=

$$\Big\{\Big(\left(2\,L2^2 + 3\,x0^2 + 6\,b\,\sigma^2\right)\Big(4\,L2^4\,(p1 - p2) + 2\,L2^3\,(-3 + 2\,p0 - p1 - p2)\,x0 + 3\,L2\,(4 + 2\,p0 - p1 - p2)\,x0^3 + 24\,b\,L2^2\,(p1 - p2)\,\sigma^2 + 6\,b\,L2\,(-4 + 2\,p0 - p1 - p2)\,x0\,\sigma^2 - 9\,(p1 - p2)\left(x0^4 - 4\,b^2\,\sigma^4\right)\Big)\Big) \Big/ \Big(4\left(4\,L2^5 + 9\,L2\left(x0^2 - 2\,b\,\sigma^2\right)^2 + L2^3\left(-9\,x0^2 + 24\,b\,\sigma^2\right)\right)\Big)\Big\}$$

Replace : pt = p1 + p2 - 2 p3, d12=p1-p2. Center measurement is measurement 3.

In[ ]:=
```
rcr = FullSimplify[rc /. p0 → 1 / 2 * (p1 + p2 - pt) /. p1 → d12 + p2, L2 > 0]
```

Out[ ]=

$$\Big\{\Big(\left(2\,L2^2 + 3\,x0^2 + 6\,b\,\sigma^2\right)\Big(L2\,x0\left(-2\,L2^2\,(3 + pt) - 3\,(-4 + pt)\,x0^2 - 6\,b\,(4 + pt)\,\sigma^2\right) + d12\left(-9\,x0^4 + 4\left(L2^2 + 3\,b\,\sigma^2\right)^2\right)\Big)\Big) \Big/ \Big(4\left(4\,L2^5 + 9\,L2\left(x0^2 - 2\,b\,\sigma^2\right)^2 + L2^3\left(-9\,x0^2 + 24\,b\,\sigma^2\right)\right)\Big)\Big\}$$

In[ ]:=
```
rcr /. b → 0 // FullSimplify
```

Out[ ]=

$$\left\{\frac{\left(2\,L2^2+3\,x0^2\right)\left(4\,d12\,L2^4-2\,L2^3\,(3+pt)\,x0-3\,L2\,(-4+pt)\,x0^3-9\,d12\,x0^4\right)}{4\left(4\,L2^5-9\,L2^3\,x0^2+9\,L2\,x0^4\right)}\right\}$$

##### LSQ formulation, find minima directly, 2 point measurement.

In[ ]:=
```
pn = pnorm[f_q, {{-L2}, {L2}}] /. b → 0;
s = Solve[
   (-2 * ((p1 - pn⟦1⟧) * D[pn⟦1⟧, x0] + (p2 - pn⟦2⟧) * D[pn⟦2⟧, x0])) == 0 /. x0s → x0, {x0}];
rd2 = FullSimplify[s, Assumptions → L2 > 0]
```

Out[ ]=

$$\left\{\{x0\to -L2\},\ \{x0\to L2\},\ \left\{x0\to \frac{L2-L2\,\sqrt{1-p1^2+2\,p1\,p2-p2^2}}{p1-p2}\right\},\right.$$
$$\left.\left\{x0\to \frac{L2\left(1+\sqrt{-((-1+p1-p2)\,(1+p1-p2))}\right)}{p1-p2}\right\}\right\}$$

Solutions 1, 2 are maxima, solutions 3, 4 are minima.

In[ ]:=
```
FullSimplify[rd2⟦{3, 4}⟧ /. p1 → d12 + p2, L2 > 0]
```

Out[ ]=

$$\left\{\left\{x0\to \frac{L2-\sqrt{1-d12^2}\,L2}{d12}\right\},\ \left\{x0\to \frac{\left(1+\sqrt{1-d12^2}\right)L2}{d12}\right\}\right\}$$

---

### Direct solution setting experimental measurements equal to theoretical

##### 3 Point Measurement, background is free fitting parameter

In[ ]:=
```
eq = {f_q[-L2] == p1, f_q[L2] == p2, f_q[0] == p0}
```

Out[ ]=

$$\left\{A\,b+\frac{A\,(-L2-x0)^2}{2\,\sigma^2}==p1,\ A\,b+\frac{A\,(L2-x0)^2}{2\,\sigma^2}==p2,\ A\,b+\frac{A\,x0^2}{2\,\sigma^2}==p0\right\}$$

In[ ]:=
```
sol3 = Solve[eq, {x0, A, b}]
```

Out[ ]=

$$\left\{\left\{x0\to \frac{-L2\,p1+L2\,p2}{2\,(2\,p0-p1-p2)},\ A\to \frac{-2\,p0\,\sigma^2+p1\,\sigma^2+p2\,\sigma^2}{L2^2},\right.\right.$$
$$\left.\left.b\to -\frac{L2^2\left(16\,p0^2-8\,p0\,p1+p1^2-8\,p0\,p2-2\,p1\,p2+p2^2\right)}{8\,(2\,p0-p1-p2)^2\,\sigma^2}\right\}\right\}$$

In[∘]:=
```
FullSimplify[sol3 /. p0 → 1 / 2 * (p1 + p2 - pt) /. p1 → d12 + p2, L2 > 0]
```

Out[∘]=

$$\left\{\left\{x0 \to \frac{d12\ L2}{2\ pt},\ A \to \frac{pt\ \sigma^2}{L2^2},\ b \to -\frac{L2^2\ \left(d12^2 - 4\ d12\ pt + 4\ pt\ (-2\ p2 + pt)\right)}{8\ pt^2\ \sigma^2}\right\}\right\}$$

#### 2 Point measurement

In[∘]:=
```
eq2 = {f_q[-L2] == p1, f_q[L2] == p2} /. b → 0
```

Out[∘]=

$$\left\{\frac{A\ (-L2 - x0)^2}{2\ \sigma^2} == p1,\ \frac{A\ (L2 - x0)^2}{2\ \sigma^2} == p2\right\}$$

In[∘]:=
```
sol2s = FullSimplify[sol2 = Solve[eq2, {x0, A}], {L2 > 0, p1 > 0, p2 > 0}]
```

Out[∘]=

$$\left\{\left\{x0 \to \frac{L2\ \left(p1 + p2 + 2\ \sqrt{p1\ p2}\right)}{p1 - p2},\ A \to \frac{\left(p1 + p2 - 2\ \sqrt{p1\ p2}\right)\ \sigma^2}{2\ L2^2}\right\},\right.$$
$$\left.\left\{x0 \to L2\ \left(-1 + \frac{2\ \sqrt{p1}}{\sqrt{p1} + \sqrt{p2}}\right),\ A \to \frac{\left(p1 + p2 + 2\ \sqrt{p1\ p2}\right)\ \sigma^2}{2\ L2^2}\right\}\right\}$$

#### Compare with MLE

Direct 2 point solution is equal to MLE

In[∘]:=
```
xmle1 = -L / 2 + L / (1 + Sqrt[p2] / Sqrt[p1]);
xmle2 = -L / 2 + L / (1 - Sqrt[p2] / Sqrt[p1]);
FullSimplify[xmle2 - x0 /. sol2s⟦1⟧ /. L2 → L / 2, {p1 > 0, p2 > 0}]
```

Out[∘]=

0

### LSQ for 2 D donut, quadratic approximation

Background set to zero, can be subtracted from measurements. We define pattern positions.

In[]:=
```
fD[x_, y_] := A ((x - x0)^2 + (y - y0)^2)/(2 σ^2)
R = L / 2;
M3c = {{ang2x[0 π, R], ang2y[0 π, R]},
   {ang2x[2 π / 3, R], ang2y[2 π / 3, R]}, {ang2x[4 π / 3, R], ang2y[4 π / 3, R]}, {0, 0}};
M3 = {{ang2x[0 π, R], ang2y[0 π, R]},
   {ang2x[2 π / 3, R], ang2y[2 π / 3, R]}, {ang2x[4 π / 3, R], ang2y[4 π / 3, R]}};
M4c = {{ang2x[0 π, R], ang2y[0 π, R]}, {ang2x[2 π / 4, R], ang2y[2 π / 4, R]},
   {ang2x[4 π / 4, R], ang2y[4 π / 4, R]}, {ang2x[6 π / 4, R], ang2y[6 π / 4, R]}, {0, 0}};
M4 = {{ang2x[0 π, R], ang2y[0 π, R]}, {ang2x[2 π / 4, R], ang2y[2 π / 4, R]},
   {ang2x[4 π / 4, R], ang2y[4 π / 4, R]}, {ang2x[6 π / 4, R], ang2y[6 π / 4, R]}};
M6c = {{ang2x[0 π, R], ang2y[0 π, R]}, {ang2x[2 π / 6, R], ang2y[2 π / 6, R]},
   {ang2x[4 π / 6, R], ang2y[4 π / 6, R]}, {ang2x[6 π / 6, R], ang2y[6 π / 6, R]},
   {ang2x[8 π / 6, R], ang2y[8 π / 6, R]}, {ang2x[10 π / 6, R], ang2y[10 π / 6, R]}, {0, 0}};
M6 = {{ang2x[0 π, R], ang2y[0 π, R]}, {ang2x[2 π / 6, R], ang2y[2 π / 6, R]},
   {ang2x[4 π / 6, R], ang2y[4 π / 6, R]}, {ang2x[6 π / 6, R], ang2y[6 π / 6, R]},
   {ang2x[8 π / 6, R], ang2y[8 π / 6, R]}, {ang2x[10 π / 6, R], ang2y[10 π / 6, R]}};
```

#### LSQ for 2D donut around x0==0:

Here we follow Eilers's section 2.2 on page 34. 3 orbit points, no center

In[]:=
```
pn = pnorm[fD, M3c];
J = FullSimplify[D[pn, {{x0, y0}}] /. x0 → 0 /. y0 → 0];
pv = {p1, p2, p3, p0};
Y = pv - pn /. x0 → 0 /. y0 → 0 // Simplify;
r = lsq[J, Y]
```

Out[]=

$$\left\{-\frac{1}{4} L (2\, p1 - p2 - p3),\ -\frac{1}{4} \sqrt{3}\, L (p2 - p3)\right\}$$

In[]:=
```
-Total[pv * M3c] // FullSimplify
```

Out[]=

$$\left\{\frac{1}{4} L (-2\, p1 + p2 + p3),\ \frac{1}{4} \sqrt{3}\, L (-p2 + p3)\right\}$$

That means: rest = - p·rpattern

#### Around x0 for iterative LSQ, 4 orbit points + center

In[∘]:=
```
pn = pnorm[f_D, M4c];
J = FullSimplify[D[pn, {{x0, y0}}]];
pv = {p1, p2, p3, p4, p0};
Y = pv - pn // Simplify;
dr = lsq[J, Y];
dr // FullSimplify
```

Out[∘]=

$$\left\{-\frac{1}{4 L^5-30 L^3 \left(x0^2+y0^2\right)+100 L \left(x0^2+y0^2\right)^2} \left(L^2+5 \left(x0^2+y0^2\right)\right) \left(2 L^4 (p1-p3)+L^3 (3-4 p0+p1+p2+p3+p4)\, x0-15 L^2 y0 (-p2\, x0+p4\, x0+p1\, y0-p3\, y0)-5 L (4+4 p0-p1-p2-p3-p4)\, x0 \left(x0^2+y0^2\right)-50 \left(x0^2+y0^2\right) \left((p1-p3)\, x0^2+2 (p2-p4)\, x0\, y0+(-p1+p3)\, y0^2\right)\right),\right.$$

$$\left.-\frac{1}{4 L^5-30 L^3 \left(x0^2+y0^2\right)+100 L \left(x0^2+y0^2\right)^2} \left(L^2+5 \left(x0^2+y0^2\right)\right) \left(2 L^4 (p2-p4)+L^3 (3-4 p0+p1+p2+p3+p4)\, y0+15 L^2 x0 (-p2\, x0+p4\, x0+p1\, y0-p3\, y0)-5 L (4+4 p0-p1-p2-p3-p4)\, y0 \left(x0^2+y0^2\right)+50 \left(x0^2+y0^2\right) \left((p2-p4)\, x0^2+2 (-p1+p3)\, x0\, y0+(-p2+p4)\, y0^2\right)\right)\right\}$$

In[∘]:=
```
drs = FullSimplify[dr /. p0 → 1 / 4 * (p1 + p2 + p3 + p4 + ps) /. p2 → p4 + d24 /. p1 → p3 + d13,
   L > 0] // InputForm
```

Out[∘]//InputForm=
```
{-(((L^2 + 5*(x0^2 + y0^2))*(-(x0*(L^3*(-3 + ps) - 15*d24*L^2*y0 + 5*L*(4 + ps)*(x0^2 + y0
   (4*L^5 - 30*L^3*(x0^2 + y0^2) + 100*L*(x0^2 + y0^2)^2)),
 -(((L^2 + 5*(x0^2 + y0^2))*(-(y0*(L^3*(-3 + ps) - 15*d13*L^2*x0 + 5*L*(4 + ps)*(x0^2 + y0
   (4*L^5 - 30*L^3*(x0^2 + y0^2) + 100*L*(x0^2 + y0^2)^2))}
```

#### Around x0 for iterative LSQ, 4 orbit points

In[•]:=
```
pn = pnorm[fD, M4];
J = FullSimplify[D[pn, {{x0, y0}}]];
pv = {p1, p2, p3, p4};
Y = pv - pn // Simplify;
dr = lsq[J, Y];
dr // FullSimplify
```

Out[•]=

$$\left\{-\frac{1}{2\left(L^3-4L\left(x0^2+y0^2\right)\right)}\left(L^2\,(p1-p3)+2\,L\,x0+4\,(p1-p3)\,x0^2+8\,(p2-p4)\,x0\,y0+4\,(-p1+p3)\,y0^2\right)\left(L^2+4\left(x0^2+y0^2\right)\right),\ -\frac{\left((p2-p4)\left(L^2-4\,x0^2\right)+2\,(L+4\,(p1-p3)\,x0)\,y0+4\,(p2-p4)\,y0^2\right)\left(L^2+4\left(x0^2+y0^2\right)\right)}{2\left(L^3-4L\left(x0^2+y0^2\right)\right)}\right\}$$

In[•]:=
```
drs = FullSimplify[dr /. p2 → p4 + d24 /. p1 → p3 + d13, L > 0] // InputForm
```

Out[•]//InputForm=
```
{-1/2*((L^2 + 4*(x0^2 + y0^2))*(2*x0*(L + 4*d24*y0) + d13*(L^2 + 4*(x0 - y0)*(x0 + y0))))/
 -1/2*((L^2 + 4*(x0^2 + y0^2))*(2*(L + 4*d13*x0)*y0 + d24*(L^2 - 4*x0^2 + 4*y0^2)))/(L^3 -
```

#### Around x0 for iterative LSQ, 6 orbit points + center

In[•]:=
```
pn = pnorm[fD, M6c];
J = FullSimplify[D[pn, {{x0, y0}}]];
pv = {p1, p2, p3, p4, p5, p6, p0};
Y = pv - pn // Simplify;
dr = lsq[J, Y];
dr // FullSimplify // InputForm
```

Out[•]//InputForm=
```
{-1/12*((3*L^2 + 14*(x0^2 + y0^2))*(-196*(2*p1 + p2 - p3 - 2*p4 - p5 + p6)*x0^4 + 28*x0^3*
     2*x0*(3*L^3*(5 - 6*p0 + p1 + p2 + p3 + p4 + p5 + p6) + 35*Sqrt[3]*L^2*(p2 + p3 - p5 -
     (2*p1 + p2 - p3 - 2*p4 - p5 + p6)*(9*L^4 - 70*L^2*y0^2 + 196*y0^4)))/(9*L^5 - 70*L^3*
 -1/12*((3*L^2 + 14*(x0^2 + y0^2))*(9*Sqrt[3]*L^4*(p2 + p3 - p5 - p6) + 6*L^3*(5 - 6*p0 +
     70*L^2*x0*(Sqrt[3]*(-p2 - p3 + p5 + p6)*x0 + (2*p1 + p2 - p3 - 2*p4 - p5 + p6)*y0) +
     196*(Sqrt[3]*(p2 + p3 - p5 - p6)*x0^4 + 2*(-2*p1 - p2 + p3 + 2*p4 + p5 - p6)*x0^3*y0
   (9*L^5 - 70*L^3*(x0^2 + y0^2) + 196*L*(x0^2 + y0^2)^2)}
```

#### Around x0 for iterative LSQ, 6 orbit points no center

In[∘]:=
```
pn = pnorm[fD, M6];
J = FullSimplify[D[pn, {{x0, y0}}]];
pv = {p1, p2, p3, p4, p5, p6};
Y = pv - pn // Simplify;
dr = lsq[J, Y];
dr // FullSimplify // InputForm
```

Out[∘]//InputForm=
```
{-1/4*((4*(2*p1 + p2 - p3 - 2*p4 - p5 + p6)*x0^2 + 4*x0*(L + 2*Sqrt[3]*(p2 + p3 - p5 - p6)
   (L^3 - 4*L*(x0^2 + y0^2)), -1/4*(Sqrt[3]*(p2 + p3 - p5 - p6)*(L^4 - 16*x0^4) + 4*(L + 2
    16*(L + 2*(2*p1 + p2 - p3 - 2*p4 - p5 + p6)*x0)*y0^3 + 16*Sqrt[3]*(p2 + p3 - p5 - p6)*
```

#### Around x0 for iterative LSQ, 3 orbit points + center

In[∘]:=
```
pn = pnorm[fD, M3c];
J = FullSimplify[D[pn, {{x0, y0}}]];
pv = {p1, p2, p3, p0};
Y = pv - pn // Simplify;
dr = lsq[J, Y];
dr // FullSimplify // InputForm
```

Out[∘]//InputForm=
```
{-1/12*((3*L^2 + 16*(x0^2 + y0^2))*(9*L^4*(2*p1 - p2 - p3) + 12*L^3*(2 - 3*p0 + p1 + p2 +
      64*L*(3 + 3*p0 - p1 - p2 - p3)*x0*(x0^2 + y0^2) - 256*(x0^2 + y0^2)*((2*p1 - p2 - p3)
   (9*L^5 - 64*L^3*(x0^2 + y0^2) + 256*L*(x0^2 + y0^2)^2),
 -1/12*((3*L^2 + 16*(x0^2 + y0^2))*(9*Sqrt[3]*L^4*(p2 - p3) + 12*L^3*(2 - 3*p0 + p1 + p2 +
      256*(Sqrt[3]*(p2 - p3)*x0^4 + 2*(-2*p1 + p2 + p3)*x0^3*y0 + 2*(-2*p1 + p2 + p3)*x0*y0
   (9*L^5 - 64*L^3*(x0^2 + y0^2) + 256*L*(x0^2 + y0^2)^2)}
```

#### Around x0 for iterative LSQ, 3 orbit points no center

In[∘]:=
```
pn = pnorm[fD, M3];
J = FullSimplify[D[pn, {{x0, y0}}]];
pv = {p1, p2, p3};
Y = pv - pn // Simplify;
dr = lsq[J, Y];
dr // FullSimplify // InputForm
```

Out[∘]//InputForm=
```
{-1/4*((4*(2*p1 - p2 - p3)*x0^2 + 4*x0*(L + 2*Sqrt[3]*(p2 - p3)*y0) + (2*p1 - p2 - p3)*(L^
 -1/4*((L^2 + 4*(x0^2 + y0^2))*(Sqrt[3]*L^2*(p2 - p3) + 4*L*y0 - 4*(Sqrt[3]*(p2 - p3)*x0^2
```

### LSQ for 2 D Gauss, linear approx. around x=0

#### Center measurement, N=4

xest = (no + Exp[Ls2])/(Ls2 no)* Sum[$p_i \cdot$ rpat$_i$ ], Ls2 = $L^2/8\,\sigma^2$ for center measurements

In[ ]:=
```
f_G[x_, y_] := A Exp[-((x - x0)^2 + (y - y0)^2)/(2 σ^2)];
pn = pnorm[f_G, M4c];
J = FullSimplify[D[pn, {{x0, y0}}] /. x0 → 0 /. y0 → 0];
pv = {p1, p2, p3, p4, p0};
Y = pv - pn /. x0 → 0 /. y0 → 0 // Simplify;
r = lsq[J, Y]
Ls2 = L^2 / 8 / σ^2;
xesteq = (4 + Exp[Ls2]) / Ls2 / 4 * Total[pv * M4c] // FullSimplify;
xesteq - r // Simplify
```

Out[ ]=
$$\left\{\frac{\left(4+e^{\frac{L^2}{8\sigma^2}}\right)(p1-p3)\,\sigma^2}{L},\ \frac{\left(4+e^{\frac{L^2}{8\sigma^2}}\right)(p2-p4)\,\sigma^2}{L}\right\}$$

Out[ ]=
{0, 0}

#### Center measurement, N = 6

In[ ]:=
```
pn = pnorm[f_G, M6c];
J = FullSimplify[D[pn, {{x0, y0}}] /. x0 → 0 /. y0 → 0];
pv = {p1, p2, p3, p4, p5, p6, p0};
Y = pv - pn /. x0 → 0 /. y0 → 0 // Simplify;
r = lsq[J, Y]
Ls2 = L^2 / 8 / σ^2;
xesteq = (6 + Exp[Ls2]) / Ls2 / 6 * Total[pv * M6c] // FullSimplify;
xesteq - r // Simplify
```

Out[ ]=
$$\left\{\frac{\left(6+e^{\frac{L^2}{8\sigma^2}}\right)(2\,p1+p2-p3-2\,p4-p5+p6)\,\sigma^2}{3\,L},\ \frac{\left(6+e^{\frac{L^2}{8\sigma^2}}\right)(p2+p3-p5-p6)\,\sigma^2}{\sqrt{3}\,L}\right\}$$

Out[ ]=
{0, 0}

#### No center measurement, N=4

xest = 1 / (Ls2) * Sum[$p_i \cdot$ rpat$_i$ ], Ls2 = L ^ 2 / 8 / σ ^ 2

In[∘]:=
```
pn = pnorm[f_G, M4];
J = FullSimplify[D[pn, {{x0, y0}}] /. x0 → 0 /. y0 → 0];
pv = {p1, p2, p3, p4};
Y = pv - pn /. x0 → 0 /. y0 → 0 // Simplify;
r = lsq[J, Y]
ph = {p1, p2, p3, p4};
xesteq = Total[ph * M4] / Ls2 // FullSimplify;
xesteq - r // Simplify
```

Out[∘]=

$$\left\{\frac{4\ (p1 - p3)\ \sigma^2}{L}, \frac{4\ (p2 - p4)\ \sigma^2}{L}\right\}$$

Out[∘]=

{0, 0}

### LSQ for 2 D donut, linear approx. around x=0

#### No measurement, N=4

In[∘]:=

$$f_{do}[x\_, y\_] := A \frac{(x - x0)^2 + (y - y0)^2}{4\ \pi\ \sigma^4} \text{Exp}\left[-\frac{(x - x0)^2 + (y - y0)^2}{2\ \sigma^2}\right];$$

```
pn = pnorm[f_do, M4c];
J = FullSimplify[D[pn, {{x0, y0}}] /. x0 → 0 /. y0 → 0];
pv = {p1, p2, p3, p4, p0};
Y = pv - pn /. x0 → 0 /. y0 → 0;
r = lsq[J, Y] // FullSimplify
xesteq = 1 / (L^2 / (8 σ^2) - 1) * Total[pv * M4c] // FullSimplify;
xesteq - r // Simplify
```

Out[∘]=

$$\left\{\frac{4\ L\ (p1 - p3)\ \sigma^2}{L^2 - 8\ \sigma^2}, \frac{4\ L\ (p2 - p4)\ \sigma^2}{L^2 - 8\ \sigma^2}\right\}$$

Out[∘]=

{0, 0}

#### No measurement, N=4

In[∘]:=
$$f_B[x\_, y\_] := A\, 4\, \text{Exp}[1]\, \text{Log}[2]\, \frac{(x - x0)^2 + (y - y0)^2}{\text{fwhm}^2}\, \text{Exp}\left[-4\, \text{Log}[2]\, \frac{(x - x0)^2 + (y - y0)^2}{\text{fwhm}^2}\right];$$

```
pn = pnorm[fB, M4c];
J = FullSimplify[D[pn, {{x0, y0}}] /. x0 → 0 /. y0 → 0];
pv = {p1, p2, p3, p4, p0};
Y = pv - pn /. x0 → 0 /. y0 → 0;
r = lsq[J, Y] // FullSimplify
xesteq = 1 / (Log[2 ] L^2 / (fwhm^2) - 1) * Total[pv * M4c] // FullSimplify;
xesteq - r // Simplify
```

Out[∘]=
$$\left\{\frac{\text{fwhm}^2\, L\, (p1 - p3)}{-2\, \text{fwhm}^2 + L^2\, \text{Log}[4]}, \frac{\text{fwhm}^2\, L\, (p2 - p4)}{-2\, \text{fwhm}^2 + L^2\, \text{Log}[4]}\right\}$$

Out[∘]=
{0, 0}

In[∘]:=

#### Quadratic approximation of donut

In[∘]:=
$$f_r[r\_] := A\, \frac{(r)^2}{4\, \pi\, \sigma^4}\, \text{Exp}\left[-\frac{r^2}{2\, \sigma^2}\right];$$

In[∘]:=
$$\text{Series}[f_r[x], \{x, 0, 2\}]$$

Out[∘]=
$$\frac{A\, x^2}{4\, \pi\, \sigma^4} + O[x]^3$$
